# Reliable delineation of *Clostridioides difficile* and related members of the family Peptostreptococcaceae using phylogenomics and spore coat protein–specific molecular markers

**DOI:** 10.64898/2025.12.05.692649

**Authors:** Jianying Han, Yannan Li, Yuxi Xu, Ji Zeng, Shaoting Li

## Abstract

Traditional bacterial classification relies on phenotypic traits (e.g., morphology, metabolic profiles), but these methods lack resolution for closely related taxa and are biased by culture conditions. While 16S rRNA gene sequencing is a widely used molecular complement, it fails to resolve closely related Peptostreptococcaceae species, including *Clostridioides difficile*. These limitations have caused family-level taxonomic confusion and ambiguous *Clostridioides* genus boundaries, hindering clinical identification of pathogenic strains and posing public health risks. To address these limitations, we developed an integrated approach combining multi-scale phylogenomic and protein-based molecular evidence, adopting a hierarchical workflow: first, constructing a 16S rRNA phylogeny of 151 Firmicutes strains to demonstrate traditional marker inadequacies; second, generating a whole-genome protein phylogeny of 51 representative Peptostreptococcaceae genomes and defining taxonomic boundaries via Average Amino Acid Identity (AAI); third, analyzing spore-associated protein patterns across *C. difficile* isolates and related genomes. Results revealed high conservation of *C. difficile* spore coat/exosporium proteins and clear genus-level phylogenetic distinctiveness of these proteins. Combined with AAI-validated whole-genome data, our findings support key Peptostreptococcaceae taxonomic revisions: redefining polyphyletic *Romboutsia*, reassigning *Eubacterium tenue* to *Paeniclostridium*, and elevating *Alkalithermobacter* to genus status. This study establishes spore coat proteins as core taxonomic markers for spore-forming bacteria, with our integrated strategy overcoming traditional limitations to improve classification accuracy and *C. difficile* surveillance.

**Importance:** Conventional classification struggles to resolve closely related Peptostreptococcaceae species (e.g., *C. difficile*). We developed an integrated framework combining 16S rRNA sequencing, whole-genome protein analysis, and spore trait assessment, with a key innovation: identifying spore coat/exosporium proteins as robust, conserved taxonomic markers. This approach enabled three pivotal Peptostreptococcaceae revisions—redefining *Romboutsia*, reassigning *Eubacterium tenue* to *Paeniclostridium*, and elevating *Alkalithermobacter* to genus rank. The findings resolve a longstanding microbial systematics bottleneck for spore-forming bacteria, provide critical taxonomic context for *C. difficile*’s precise monitoring and prevention, and expand taxonomic markers beyond nucleic acid-based methods. This advances classification precision, critical for microbial ecology, pathogenesis, and industrial microbiology research.

## Introduction

Accurate classification of bacterial species and genera is fundamental to microbiological research, enabling critical insights into microbial ecology and function (1), with direct implications for human health (2). Nowhere is this more evident than in clinical medicine (2), where precise taxonomic assignment of pathogens is indispensable for elucidating mechanisms of virulence and developing targeted interventions (3). This necessity is starkly illustrated by *Clostridioides difficile* (*C. difficile*), a leading cause of antibiotic-associated diarrhea globally. *C. difficile* infection presents a spectrum of disease, from mild diarrhea to life-threatening pseudomembranous colitis, and is notably prone to recurrence (4), incurring healthcare costs amounting to billions of dollars annually (5, 6). Concerns are amplified in regions like China, where high toxin detection rates in tertiary hospitals suggest a potentially underestimated epidemic burden (7, 8). These clinical realities underscore the imperative to resolve the precise phylogenetic position and and taxonomic boundaries of *C. difficile*.

The taxonomic home of *C. difficile* lies within the family Peptostreptococcaceae, a group whose classification requires rigorous phylogenetic analysis. *C. difficile* itself was reclassified from the genus *Clostridium* to *Clostridioides* following genomic evidence (9). Peptostreptococcaceae species, which inhabit diverse niches including the human gut and soil, often share key biological traits with *C. difficile*, such as anaerobic metabolism and sporulation (10). However, the current taxonomy of this family is fraught with challenges that obscure phylogenetic relationships (11). Traditional phenotypic methods lack the resolution to distinguish between closely related genera that share these common traits (12), and their results can be biased by culture conditions, ultimately compromising the accuracy of taxonomic assignments and leaving the placement of many strains ambiguous (10, 13).

While molecular sequencing promised to overcome these limitations, both 16S rRNA and whole-genome approaches exhibit specific shortcomings that perpetuate taxonomic controversies (14). The 16S rRNA gene, a cornerstone of bacterial taxonomy, is too conserved for reliable species-level discrimination. Within Firmicutes, this is exemplified by near-identical 16S sequences (>99.5% similarity) between *Bacillus subtilis* and *B. amyloliquefaciens* (15) and minimal differences between *C. difficile* and close relatives like *Clostridium sordellii* (16). Whole-genome sequencing, though more powerful, provides a definitive resolution in such cases, as demonstrated by its role in underpinning the reclassification of *C. difficile*, but it introduces challenges of cost, analytical complexity, and the difficulty of extracting phylogenetically robust signals from massive datasets (17).

These technical limitations are concretely reflected in the persistent taxonomic ambiguities within Peptostreptococcaceae. A critical issue is the non-monophyly of some described genera (12). The genus *Romboutsia*, for example, is not a natural group; some of its species show a closer phylogenetic affinity to *Paraclostridium* than to other *Romboutsia* members, blurring the evolutionary picture around *C. difficile* (10). Similarly, the species *Eubacterium tenue* displays ambiguous clustering in phylogenetic trees, sometimes associating with *Paeniclostridium* and other times with *Clostridioides*, which hinders a clear understanding of its relationship to *C. difficile*. These specific cases underscore that accurate phylogenetic reconstruction of the entire family remains an unresolved challenge, one that directly impedes the clarification of *C. difficile*’s closest relatives.

Given the constraints of sequence-based methods, functional biomarkers offer a compelling complementary strategy for taxonomic delineation. The utility of this approach is demonstrated by toxin-based classification systems for *C. difficile*, which effectively leverage the direct link between a key phenotypic trait and taxonomic identity. Beyond toxins, spore coat proteins present a promising yet underexplored class of biomarkers. Unique to spore-forming Firmicutes, these proteins possess a dual utility: their sequences harbor phylogenetic signal (18), while their functions are directly tied to environmental persistence and pathogenicity. Alterations in *C. difficile* spore coat proteins, for instance, can impact intestinal colonization and disease progression. Despite this potential, a comprehensive, spore coat protein-based taxonomic framework for the Peptostreptococcaceae remains to be established, and its power to resolve phylogenetic disputes requires systematic validation.

To address these gaps, our study employs an integrated strategy that combines genomic data with the analysis of characteristic protein markers. We aim to resolve the taxonomic ambiguities within the Peptostreptococcaceae family, thereby clarifying the phylogenetic context of *C. difficile* and refining the boundaries of the genus *Clostridioides*. Our methodology proceeds in three concerted steps: first, to establish a robust phylogenetic framework using 16S rRNA and whole-genome sequences; second, to investigate the phylogeny of *C. difficile* spore coat proteins through systematic analysis of their conservation, diversity, and homology; and finally, to validate the resulting classification model using Average Amino Acid Identity (AAI). This multi-faceted approach is designed to yield a reliable taxonomy that will support future ecological studies and the precise clinical identification of pathogens within this critical family.

## METHODS

### 2.1 Genomic Data Collection and Preprocessing

First, genomic sequences related to the family Peptostreptococcaceae were retrieved from the public genomic database NCBI RefSeq (19). To ensure the representativeness of this study, 51 complete genomes were downloaded. Among these, 2 genomes corresponded to *Faecalimicrobium dakarense* FF1 and *Maledivibacter halophilus* M1, which were used as outgroups for constructing the whole-genome phylogenetic tree. The remaining 49 genomes included various bacteria isolated from human or animal tissues (e.g., intestine/feces, oral cavity, and blood) and represented multiple genera, namely *Acetoanaerobium*, *Alkalithermobacter*, *Asaccharospora*, *Clostridioides*, *Clostridium*, *Criibacterium*, *Eubacterium*, *Filifactor*, *Intestinibacter*, *Lachnospira*, *Mediannikoviicoccus*, *Metaclostridioides*, *Paeniclostridium*, *Paraclostridium*, *Paramaledivibacter*, *Peptacetobacter*, *Peptoanaerobacter*, *Peptoclostridium*, *Peptostreptococcus*, *Proteocatella*, *Romboutsia*, *Tepidibacter*, *Terrisporobacter*, and *Wukongibacter*.

The selection criteria for the genomic sequences were as follows: (1) all sequences were complete genomes; (2) the sources covered multiple host types (e.g., humans and animals) to fully reflect the diversity of the Peptostreptococcaceae family across different ecological environments; (3) all genomes had clear gene annotations and functional annotations, ensuring the reliability of subsequent analyses.

Additionally, genomic data of *C. difficile* were obtained from the NCBI SRA database (20). In this study, 280 genomic sequences of *C. difficile* strains isolated from the United States were retrieved, covering 23 distinct PCR ribotypes from both clinical and environmental origins. These PCR ribotypes included PCR ribotype 027 (a hypervirulent epidemic strain of *C. difficile*), PCR ribotype 078 (a zoonosis-related strain), PCR ribotype 106, and PCR ribotype 147. These data were primarily used for systematic analysis of the presence, sequence conservation variations, and genomic distribution characteristics of key sporulation genes (e.g., *spoIVA*, *cotE*, *sleC*) across *C. difficile* strains. Moreover, they provided basic data support for subsequent studies on the associations among sporulation ability, strain virulence, and environmental adaptability.

All genomic data underwent quality control via a standardized preprocessing pipeline, with specific steps and tools detailed below. First, raw sequencing data quality assessment was performed using FastQC v0.11.9 (21).Subsequently, low-quality sequences were filtered using Trimmomatic v0.39 (22) with the parameters: ILLUMINACLIP:TruSeq3-PE.fa:2:30:10 LEADING:3 TRAILING:3

SLIDINGWINDOW:4:15 MINLEN:36. These parameters removed adapter sequences, terminal bases with a quality score < 3, and fragments < 36 bp. Next, redundant data were further cleaned using CD-HIT v4.8.1 (23) by setting a 95% similarity threshold (-c 0.95) and 90% coverage thresholds for longer (-aL 0.9) and shorter (-aS 0.9) sequences to eliminate duplicate and highly similar sequences.

Finally, gene annotation and functional prediction were conducted using Prokka (24). Default parameters were supplemented with manual correction to focus on extracting structural and functional information of spore formation-related genes, ensuring the reliability of subsequent analyses.

### 2.2 Analysis of 16S rRNA Gene Sequences

To construct a 16S rRNA-based phylogenetic tree and validate the taxonomic rationality of strains within the phylum Firmicutes, this study first established the screening criteria and sources for 16S rRNA gene sequences. A total of 151 bacterial 16S rRNA gene sequences were downloaded from the NCBI GenBank database (25), selected based on multiple peer-reviewed studies and stringent quality filters. These sequences met the following 16S rRNA-specific quality requirements: sequence integrity ≥ 95%, base call error rate < 0.1%, and annotation completeness ≥ 90%. Importantly, while the Firmicutes phylum comprises substantially more than 151 strains, the final 151 sequences included in this analysis were determined by integrating literature-derived evidence with the aforementioned quality benchmarks. Additionally, the corresponding strains were sourced from typical Firmicutes niches (e.g., human gut, animal gut, soil, water, and clinical specimens) to ensure representative sampling.

The 151 sequences corresponded to strains belonging to 16 identified families, 1 unidentified family, and 48 genera, covering most known representative families and genera within Firmicutes. The 16 identified families included Peptostreptococcaceae, Bacillaceae, Alicyclobacillaceae, Peptoniphilaceae, Lachnospiraceae, Paenibacillaceae, Clostridiaceae, Cytobacillaceae, Enterococcaceae, Eubacteriaceae, Atopobiaceae, Neomoorellaceae, Tepidibacteraceae, Streptococcaceae, Thermoanaerobacteraceae, and Sporolactobacillaceae. The unidentified family was temporarily classified as an unclassified family of Firmicutes. The 48 genera included *Clostridioides* and *Paraclostridium* (from Peptostreptococcaceae), *Bacillus* and *Paenibacillus* (from *Bacillaceae*), and *Lachnospira* and *Roseburia* (from *Lachnospiraceae*), among others. See Table S1.

Maximum likelihood (ML) phylogenetic analysis was conducted following a standardized pipeline. First, multiple sequence alignment (MSA) of the 16S rRNA gene sequences was generated using MAFFT v7.490 (26) with the G-INS-i algorithm, which is well-suited for aligning highly conserved gene sequences such as 16S rRNA due to its accuracy in resolving positional homology. The optimal nucleotide substitution model was then determined using ModelFinder (27) as implemented in IQ-TREE 2 (28), with the TVMe+R10 model selected as best-fitting under the Bayesian Information Criterion (BIC). Phylogenetic tree reconstruction was subsequently performed using IQ-TREE 2 with 1000 bootstrap replicates to evaluate branch support. Finally, the resulting tree was visualized and annotated using FigTree v1.4.4.

### 2.3 Construction of Phylogenetic Tree Based on Strain Protein Sequences

Using the protein sequences of 51 Peptostreptococcaceae strains, see Table S2, conserved proteins (single-copy orthologous proteins) were first screened using OrthoFinder v2.5.4 (29) combined with Diamond v2.1.8 (30, 31). Specifically, OrthoFinder generated orthologous gene clusters (OGs) via an orthology inference workflow based on gene clustering, while Diamond performed sequence alignment using the Smith–Waterman algorithm (--more-sensitive mode, E-value = 1e-10). The Screening of protein sequences strictly followed four criteria: (1) each strain contained only one sequence (strict single copy); (2) all 51 strains were covered; (3) the coefficient of variation of sequence length was ≤ 20% (additionally, abnormal sequences with length<100 aa or> 1000 aa were excluded); (4) average amino acid identity (AAI) ≥ 70%.

Subsequently, MUSCLE v5.1 (32) was employed for multiple sequence alignment of each screened single-copy orthologous protein individually, with multi-round iterative optimization via to improve alignment accuracy in conserved regions. Based on the MSA results, Gblocks v0.91b (33) was used for trimming, whereby only conserved blocks of length ≥ 5 aa were retained, semi-conserved gaps were allowed (with a maximum gap ratio ≤ 50%), and fragments containing > 8 consecutive non-conserved sites or gap length > 10 amino acids were removed. After trimming, SeqKit v2.4.0 (34) was used to concatenate sequences by strain, by connecting the trimmed sequences of all single-copy genes of each strain end-to-end in the order of gene IDs to form a continuous long sequence. Finally, IQ-TREE v2.2.6 (28) was used to construct the phylogenetic tree; this tool automatically selected the optimal amino acid substitution model and evaluated statistical branch reliability via 1000 ultra-fast Bootstrap replicates and aLRT tests (where Bootstrap values ≥ 70% were defined as indicating reliable branches). Subsequently, tree visualization and annotation were performed using the web-based tool tvBOT (35).

### 2.4 Conservation of Spore Coat Proteins

Genomic data from 280 U.S.-isolated *C. difficile* strains were collected, see Table S3. Concurrently, 54 spore coat proteins and 21 exosporium proteins were screened from the genome-wide proteome of *C. difficile* type strain 630 (GenBank: NC_009089.1), and their conservation was analyzed.

First, BLASTp was used for homologous sequence searches against all 280 strains’ proteomes. Hits were defined as conserved spore-related proteins only if they met simultaneous criteria: Bit score ≥ 100, E-value < 1e-10, sequence identity ≥ 90%, and sequence coverage ≥ 80%.

Subsequently, the conservation degree distribution of each spore-related protein across the 280 genomes was analyzed. Here, conservation degree refers specifically to the frequency of a spore-related protein being detected as conserved (i.e., BLASTp-identified homologs) among the 280 strains. This analysis followed three steps: first, counting strains with each conserved spore-related protein; second, calculating detection rates as (positive strains / 280) × 100%; third, classifying into grades (e.g., highly conserved: ≥80% detection; moderately conserved: 50–80%; poorly conserved: <50%) to visualize prevalence differences across proteins and strains.

### 2.5 Multiple Sequence Alignment and Phylogenetic Analysis of Spore Coat Proteins

Local BLASTp searches were performed against the proteome sequences of 49 Peptostreptococcaceae strains, using 54 *C. difficile* spore coat protein sequences and 21 *C. difficile* exosporium proteins as queries. An E-value threshold of 1e-2 was set to filter random matches, and a sequence identity threshold of ≥ 60% was used to ensure homology reliability. High-confidence homologous protein matches were obtained via these screening criteria.

Based on the screening results, a Presence/Absence matrix was constructed. If a strain’s protein sequence aligned with a spore coat protein and met the E-value and identity criteria, the corresponding position in the matrix was marked as “1” (present); otherwise, it was marked as “0” (absent). The Presence/Absence matrix was used to construct a neighbor-joining (NJ) tree in MEGA X (36) with 1000 bootstrap replicates for reliability evaluation. The Jaccard coefficient was used to calculate the distance between matrix elements, ultimately generating a phylogenetic topology with confidence support.

All analyses were performed in R version 4.4.2 (37). We assessed protein distribution by generating a heatmap with ggplot2 (38) to visualize the presence of 54 spore coat proteins and 21 exosporium proteins across 49 strains, following data organization with the tidyverse suite (39). Phylogenetic trees were constructed and manipulated separately using ape (40) and treeio (41), and visualized with ggtree. All graphics were finalized using color schemes from RColorBrewer and assembled with patchwork.

### 2.6 Multiple Sequence Alignment and Phylogenetic Analysis of Core Spore Coat Proteins

Based on the presence/absence matrix of 54 spore coat proteins and 21 exosporium proteins, a strict screening process identified 3 core spore coat proteins stably present in all 49 strains. MAFFT v7.490 (26) was used to perform multiple sequence alignment on these 3 core proteins. According to protein characteristics, trimAl v1.4.rev15 (42) was used to remove extensive “-” (insertions/deletions) from both ends of the sequences and retain core conserved regions, yielding high-quality alignment results. The final alignment results were carefully checked and optimized using the Jalview visualization tool (43).

Based on the MSA results of the 3 core spore coat proteins, a standardized amino acid variation matrix was constructed. RAxML-NG software (44) was used to construct the phylogenetic tree and evaluate its reliability. The LG+G+F substitution model (45), which accounts for amino acid frequencies and rate heterogeneity accross four GAMMA categories (46), a maximum likelihood (ML) phylogenetic tree was constructed with 1000 bootstrap replicates. Finally, FigTree was used for tree visualization to present interspecific genetic relationships and branch reliability.

### 2.7 Calculation of Average Amino-Acid Identity (AAI)

To evaluate the genetic relationships among 49 genomes, the CompareM toolkit (47) was used to calculate the AAI of conserved proteins across the whole genome. First, the Prodigal tool (48) was used to predict coding genes in each genome. Protein sequences with length ≥ 100 aa were then screened and extracted. Pairwise sequence alignment was performed using BLASTp (E-value ≤ 1e-5), and orthologous gene pairs were screened using the following criteria: alignment coverage ≥ 70% and similarity ≥ 30%. The amino acid identity of each gene pair (AAIᵢ) was calculated, and the average value was taken as the final AAI (formula: AAI = (1/n)∑ⁿᵢ₌₁ AAIᵢ, where n is the total number of orthologous gene pairs). The judgment criteria were as follows: AAI ≥ 95% (49) indicated closely related taxa; AAI ≤ 70% (50) indicated distantly related taxa (potentially belonging to different genera).

### 2.8 Construction of Spore Coat Gene Co-Expression Network

Gene expression data were obtained from the STRING database (51). A set of 54 spore coat genes and 21 exosporium-associated genes was initially queried to retrieve complete expression profiles under multiple sporulation-related conditions, including different sporulation stages, nutrient deprivation, and environmental stress. Owing to limitations in the database’s gene annotation coverage, which primarily stems from the inability to detect a subset of genes in the available experimental systems presumably because of low expression levels, only 27 genes were successfully matched with valid expression data. Subsequent co-expression network analysis was therefore restricted to these 27 genes. The network was visualized using Cytoscape v3.7.2 (52), which clearly delineated co-expression relationships among these spore coat genes and provided an intuitive model for subsequent investigation into the molecular regulatory mechanisms underlying spore coat assembly.

## Results

### Phylogenetic Analysis of the 16S rRNA Gene: Preliminary Insights into the Polyphyly of the Peptostreptococcaceae Family

Circular phylogenetic tree of 151 strains within the phylum Firmicutes, constructed using 16S rRNA gene sequences. The tree was generated via the maximum likelihood method, with values at nodes representing bootstrap support values (1,000 replicates).

To expand the analysis of overall evolutionary relationships within Firmicutes and verify the universality of taxonomic controversies in Peptostreptococcaceae, this study first reconstructed a phylogenetic tree using genomic 16S rRNA sequences of 151 representative strains. These strains cover multiple genera and species of Peptostreptococcaceae, with taxonomic annotations for all species referenced from the List of Prokaryotic Names with Standing in Nomenclature (LPSN) (53) or the Genome Taxonomy Database (GTDB) (54).

Phylogenetic analysis based on 16S rRNA gene sequences revealed that the Peptostreptococcaceae family, as defined under the current taxonomic system, is a highly paraphyletic group. Its members do not cluster into a single evolutionary clade but are scattered across at least four major clades, and each of these observations further supports the paraphyletic nature of the family. Specifically, the clade containing *Terrisporobacter petrolearius* and *Terrisporobacter mayombei* (Bootstrap support: 52–58%) is closely intertwined with *Clostridium glycolicum* and *C. felsineum* (*Clostridium* species), breaking the monophyletic boundary of Peptostreptococcaceae; although *Romboutsia* species (including *R. faecis*, *R. hominis*, and *R. ilealis*) form a distinct clade (Bootstrap support: 61%), this clade still shows close relationships with other species, suggesting incomplete separation between these groups; and *Eubacterium tenue* clusters with *Eubacteriaceae* species (Bootstrap support: 44%), directly falling outside the Peptostreptococcaceae clade. Additionally, *Peptoclostridium* and *Tepidibacter*, taxa that were originally assigned to the family Peptostreptococcaceae, are distinctly separated from the aforementioned core clades in the phylogenetic tree. These results consistently demonstrate that the current taxonomy, which is based on the traditional approach of using morphological and biochemical traits, does not accurately reflect the true phylogenetic relationships of strains in this group. They further support the view that a strictly delineated Peptostreptococcaceae family should be established using whole-genome proteomic data, with the paraphyletic components reclassified into other appropriate taxonomic units.

### Genome-Wide Phylogenomics of the Full Protein Repertoire: Resolving Genus-Level Delineation

This figure shows a phylogenetic tree constructed using protein sequences, depicting the phylogenetic relationships between Peptostreptococcaceae strains and their related taxa. The branching pattern reflects evolutionary affinities among different strains, which are inferred from differences in their protein sequences, and the numbers labeled on the branches represent bootstrap support values (based on 1000 replicates). These values indicate the reliability of each branching node, with values ≥ 70% typically considered statistically robust.

To address 16S rRNA sequencing limitations (e.g., low resolution for closely related taxa) and resolve Peptostreptococcaceae’s internal phylogenetic relationships with high precision, this study analyzed whole-genome protein sequence data. Core single-copy orthologous genes were screened, their protein sequences concatenated, and a phylogenetic tree constructed via the maximum likelihood (ML) method.

As shown in the phylogenetic tree (Figure 2), three key taxonomic patterns emerge: first, species currently assigned to the genus *Romboutsia* do not form a monophyletic group but are scattered across multiple distinct clades with substantial genetic distances between them—this indicates that the current taxonomic scope of *Romboutsia* may include non-homologous species; second, all species of the genus *Terrisporobacter* cluster tightly into a monophyletic clade with high Bootstrap support (≥ 90%), confirming its status as a stable and well-defined taxon within *Peptostreptococcaceae*; third, species of the genus *Clostridioides* (including the model strain *Clostridioides difficile* DSM 1296) form an independent monophyletic clade with strong Bootstrap support (≥ 95%), which is distinctly separated from the aforementioned *Romboutsia* and *Terrisporobacter* clades, further validating its taxonomic independence as a core genus within the family. Notably, the widespread presence of highly supported nodes (with numerous bootstrap value of 100) in the tree not only validates the high reliability of the phylogenetic relationships constructed based on protein sequences but also strengthens the credibility of the aforementioned taxonomic inferences regarding *Romboutsia*, *Terrisporobacter*, and *Clostridioides*.

**Figure 1:**
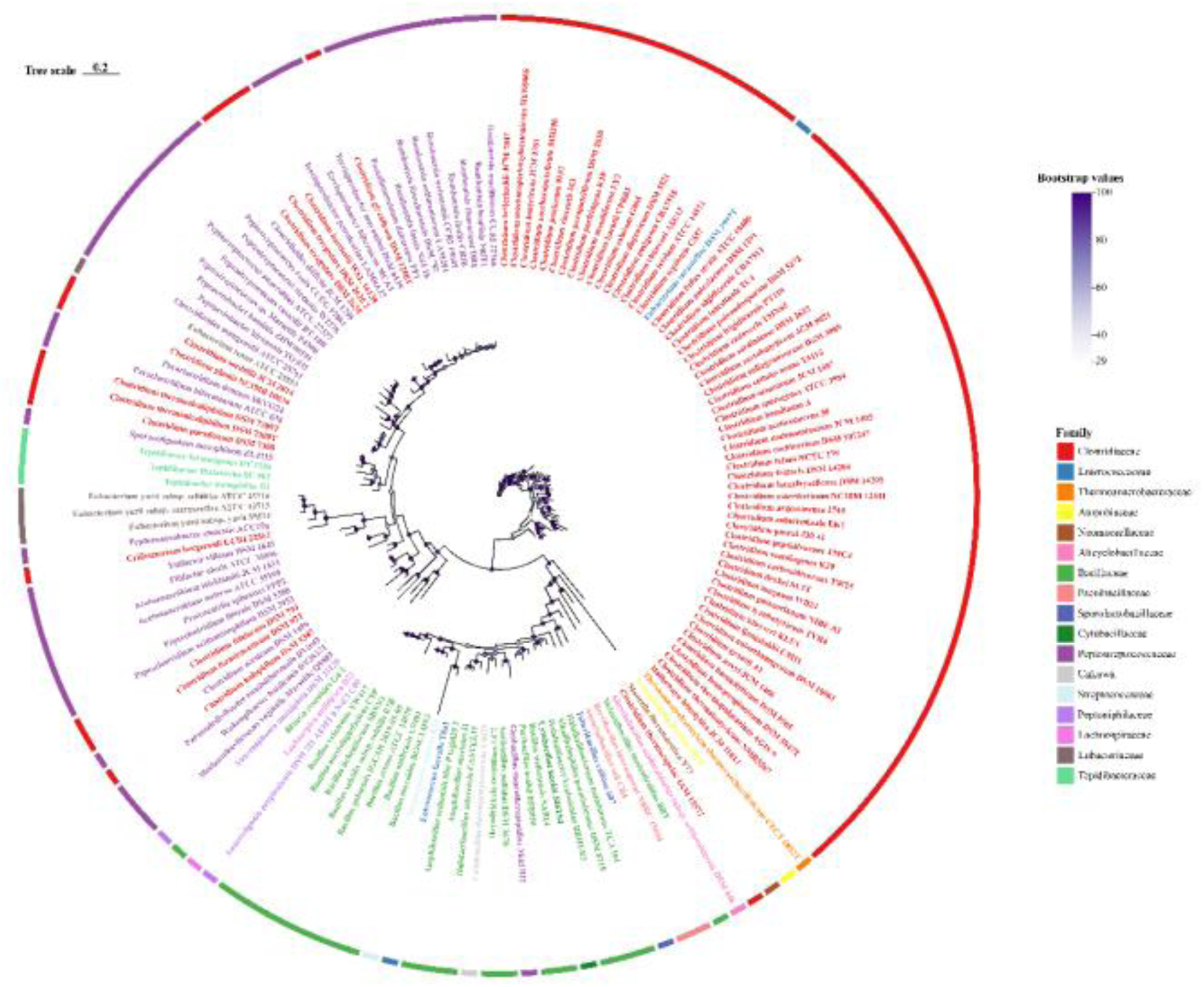
Maximum Likelihood Phylogenetic Tree Based on 16S rRNA Gene Sequences

**Figure 2:**
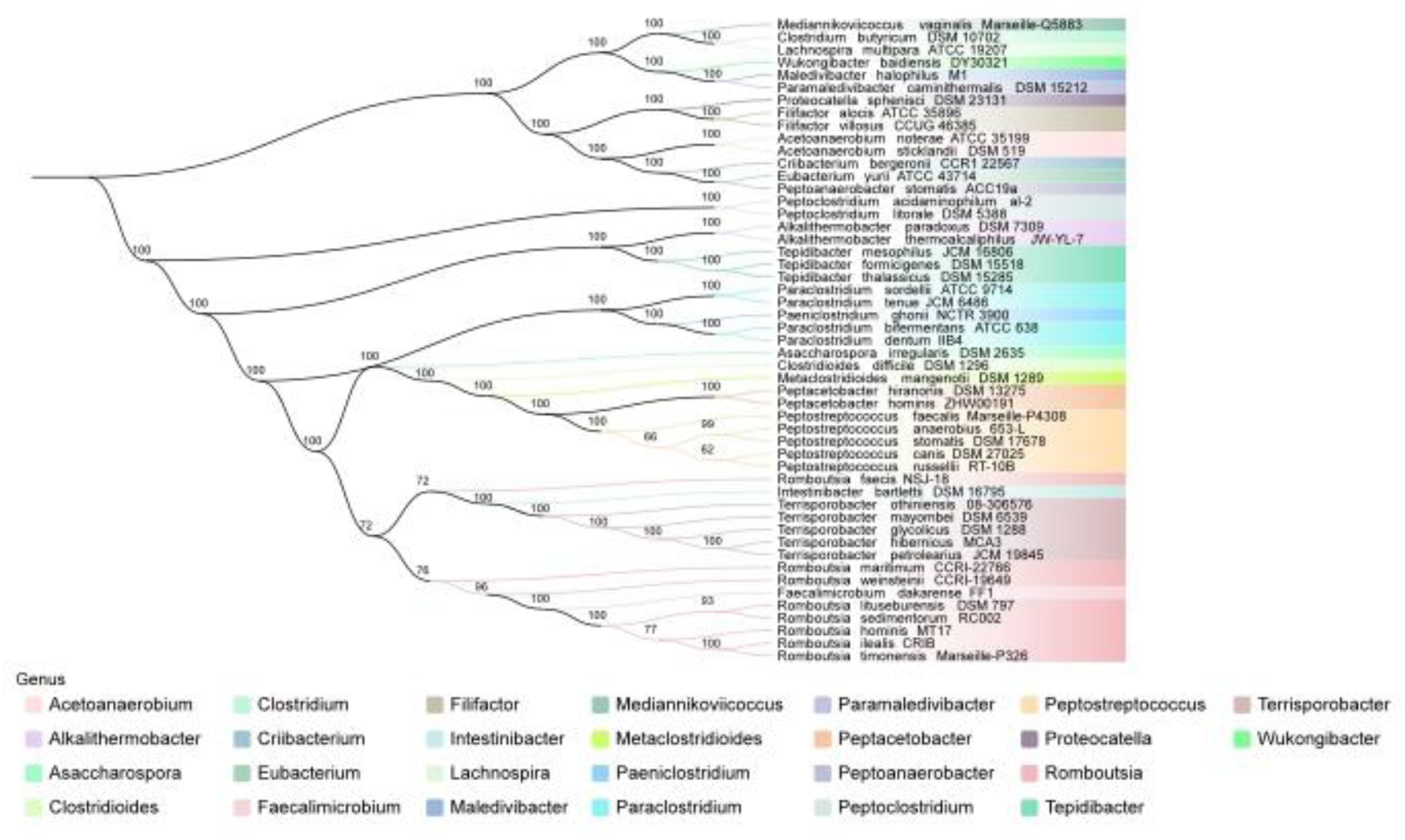
Phylogenetic Tree of Peptostreptococcaceae Strains Based on Protein Sequences

### Conservation Analysis of Spore Proteins: Supporting Their Application as Taxonomic Markers

(A) Schematic of the spore hierarchical structure. This panel illustrates the complete layered organization of the spore, including the Core (innermost region), Cortex (surrounding the core), Basement coat, Inner coat, Outer coat (collectively forming the spore coat), and Exosporium (outermost layer). The diagram also indicates the localization of spore-related proteins within these layers. (B) Conserved distribution of spore coat proteins (Basement layer and Unknown layer) in *C. difficile*. The x-axis represents the conservation rate (range: 0–1), calculated as the proportion of tested *C. difficile* strains in which a given spore coat protein was identified as conserved (defined as 100% sequence identity to the reference sequence) relative to the total number of strains. The y-axis lists individual spore coat protein names. Each color-coded bar corresponds to a protein’s conservation rate: yellow (Basement layer), green (Unknown layer). (C) Conserved distribution of exosporium proteins in *C. difficile*. Analyzed using the same method as Panel B, this panel depicts the conservation of proteins localized to the Exosporium layer. Each bar represents an individual exosporium protein, with purple indicating the Exosporium layer.

Fig. 3A illustrates the multilayered structure of the *C. difficile* spore and the distribution of spore coat proteins and exosporium proteins across distinct layers (e.g., Basement coat, Inner/Outer coat, Exosporium)—this structural context justifies analyzing protein conservation based on spatial localization. Using this framework, protein sequences from 280 clinical *C. difficile* isolates were aligned against strain 630’s reference proteome to identify spore coat proteins and exosporium proteins. Following BLASTp filtering of redundant sequences, these proteins exhibited a conservation rate of over 90% across strains (Fig. 3B); notably, 85% of these proteins showed 100% conservation (Fig. 3B, C). Collectively, spore coat proteins displayed significant sequence conservation across *C. difficile* strains, irrespective of their localization within specific layers of the spore structure. Importantly, these conserved spore coat proteins are also present in other sporulating members of the phylum Firmicutes—this cross-generic conservation suggests they may serve as novel molecular markers for phylogeny within Firmicutes, prompting further investigation of their potential utility.

**Figure 3.**
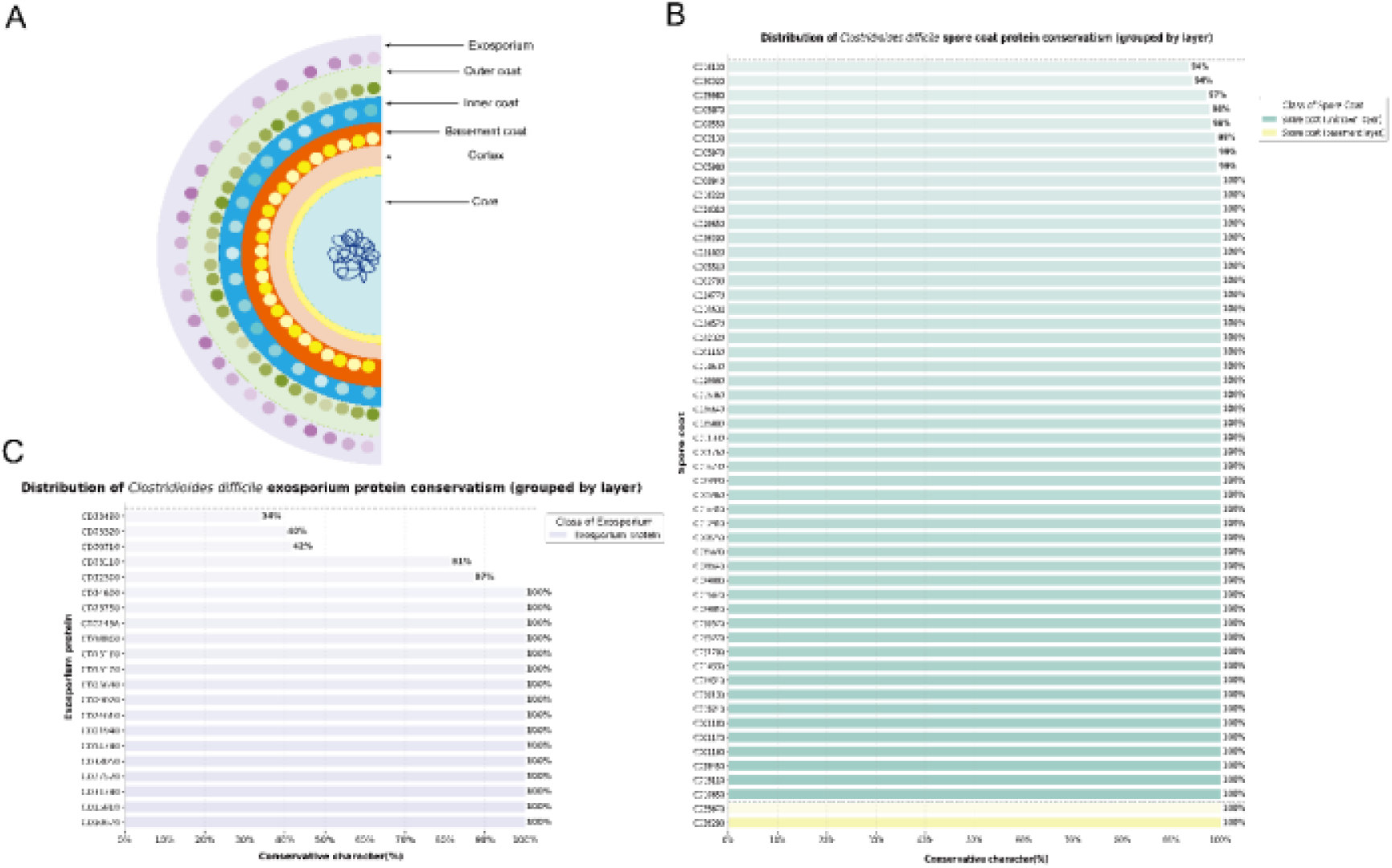
Spore layered structure and conserved distribution of spore-related proteins in *C. difficile*

### Phylogenetics of Spore Proteins: Evaluating Genus Delimitation

(A) Heatmap of BLASTp results. This heatmap presents BLASTp alignment results of strains against target proteins. On the left, a hierarchical clustering dendrogram shows strain clustering derived from sequence identity data, with clustering performed using Euclidean distance and the average linkage method. On the right, a protein similarity heatmap depicts strain-protein sequence similarity, where color intensity represents the identity percentage and dot size indicates sequence coverage. Strains are grouped by genus and color-coded for genus-level distinction. (B) Neighbor-Joining (NJ) tree based on the presence/absence matrix of 54 spore proteins and 21 exosporium proteins. This phylogenetic tree was constructed via the NJ method based on the presence/absence of 54 spore proteins and 21 exosporium proteins, with different colors corresponding to distinct genera. Supplementary GC content bar charts are included for each strain.

BLASTp alignment results showed high sequence similarity in spore formation-related proteins between *Paraclostridium* and *Metaclostridioides* strains (Fig. 4A). These proteins included key spore formation factors. Among these, SipL functions as a spore assembly protein, and SpoIVA acts as a spore cortex assembly protein. Notably, the gene clusters annotated at the bottom of the heatmap directly correspond to these proteins, a correspondence that ensures accurate identification of the target proteins. Heatmap of BLASTp results (Fig. 4A) revealed clear quantitative characteristics of sequence identity and coverage for the target proteins between the two genera. Specifically, more than 80% of the co-detected spore formation-related proteins exhibited sequence identity ≥ 80% (corresponding to the orange signal range in the heatmap) and sequence coverage ≥ 90% (corresponding to the high-coverage markers in the heatmap). This quantitative data confirm the high integrity of the conserved regions of these proteins. Collectively, these quantitative features of the heatmap directly validate high consistency in both sequence and structural integrity of the conserved spore formation-related proteins across the two genera.

**Figure 4.**
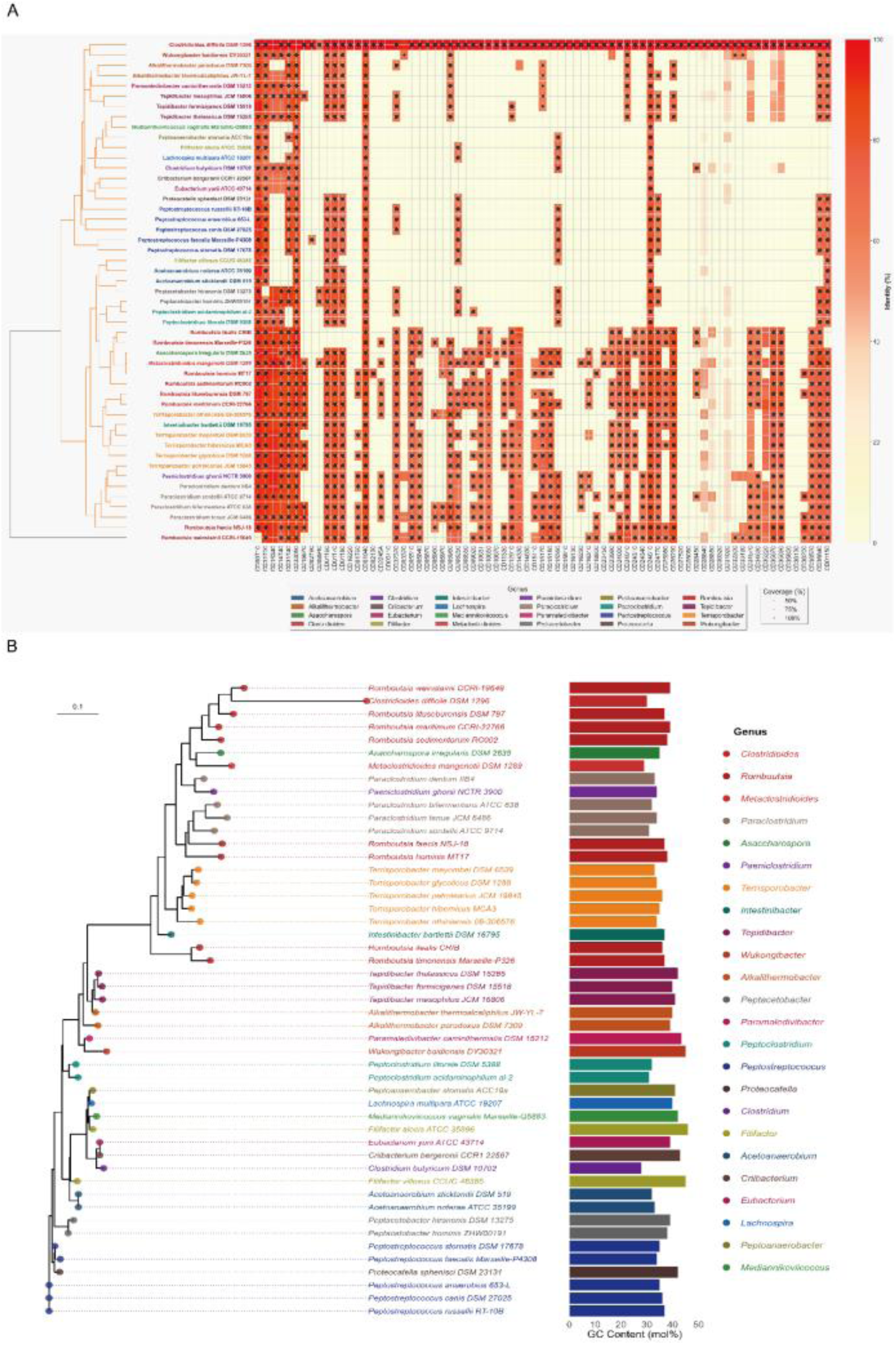
BLASTp alignment results heatmap and Neighbor-Joining phylogenetic tree based on spore/exosporium protein presence/absence matrix

Furthermore, the sequence similarity revealed by BLASTp receives strong support from phylogenetic clustering results. Specifically, *Paraclostridium* and *Metaclostridioides* cluster tightly into a single clade in the left hierarchical clustering tree. This clustering result confirms molecular consistency of the two genera in key spore formation-related conserved domains. It also reinforces their close phylogenetic relationship at the molecular level. In contrast, *Clostridioides* strains (with *Clostridioides difficile* DSM 1296 as the representative) exhibit distinct patterns that match their taxonomic divergence. The sequence similarity in the corresponding region of the heatmap for this genus exceeds 95%. This value is significantly higher than the 80% to 100% range observed for the two aforementioned genera. Its sequence coverage also exceeds 95%. Meanwhile, *Clostridioides* forms an independent clade with a distant phylogenetic relationship. These coordinated features collectively indicate substantial differences in spore-related proteins between *Clostridioides* and the two genera mentioned above (*Paraclostridium* and *Metaclostridioides*).

To prioritize the taxonomic value of spore coat proteins and exosporium proteins while avoiding sequence variation-induced biases, we first constructed a Neighbor-Joining (NJ) phylogenetic tree based on the presence-absence matrix of 54 spore coat proteins and 21 exosporium proteins (Fig. 4B). Phylogenetic analysis of this spore protein-based NJ tree yielded clear taxonomic insights: first, *Paraclostridium* and *Metaclostridioides* belong to the core phylogenetic group of the Peptostreptococcaceae family, sharing a closer phylogenetic relationship with each other than with more distantly related genera and thus forming a tight cluster; second, the genus *Clostridioides* (represented herein by *C. difficile*) exhibits an independent divergence trend, with a topology consistent with the formation of an exclusive monophyletic clade when incorporating additional conspecific strains; third, although *Romboutsia* appears monophyletic in the current tree, marked divergence among its internal subclades and close phylogenetic associations with external genera suggest a potential non-monophyletic status.

### Phylogeny of Core Spore Coat Proteins and AAI Analysis: Delineating Genetic Boundaries at the Genus Level

(A) Heatmap of Average Amino Acid Identity (AAI) results. This heatmap presents AAI values among different microbial strains, which are utilized to analyze the phylogenetic relationships and genomic similarity of the tested strains. (B) Maximum likelihood (ML) tree based on the amino acid variation matrix of three core spore coat proteins. The ML phylogenetic tree was constructed using RAxML, illustrating the evolutionary relationships among different bacterial strains. Branch colors are encoded according to the genus-level classification of strains (see the right-hand legend). Each leaf node represents a bacterial strain, with the full species name displayed as its label. Red numbers on the nodes indicate bootstrap support values (only values > 70 are shown), and branch lengths correspond to evolutionary distances.

The consistency between the topological structure and genetic distance patterns of the spore coat protein ML tree and AAI analysis (Figure 5A, B, with genera annotated by consistent color coding for improved readability) provides critical evidence for genus-level taxonomic revisions. Specifically, *Eubacterium tenue* clusters with typical *Paeniclostridium* strains (e.g., *P. ghonii*, *P. sordellii*) to form a high-confidence branch (Bootstrap support ≥ 90%), with AAI values within this branch ranging from 70% to 95%—a range that aligns with the widely accepted genetic threshold for genus-level classification (55). To further validate this threshold, we compared it with the intra-genus genetic characteristics of *Terrisporobacter* (mean AAI = 90.68, OF = 75.06): the overlapping AAI ranges between the two groups confirm that the *Eubacterium tenue-Paeniclostridium* branch’s genetic patterns are consistent with genus-level clustering. This key finding directly supports the taxonomic revision of merging *Eubacterium tenue* into *Paeniclostridium*, resolving long-standing ambiguities in their taxonomic boundaries.

**Figure 5.**
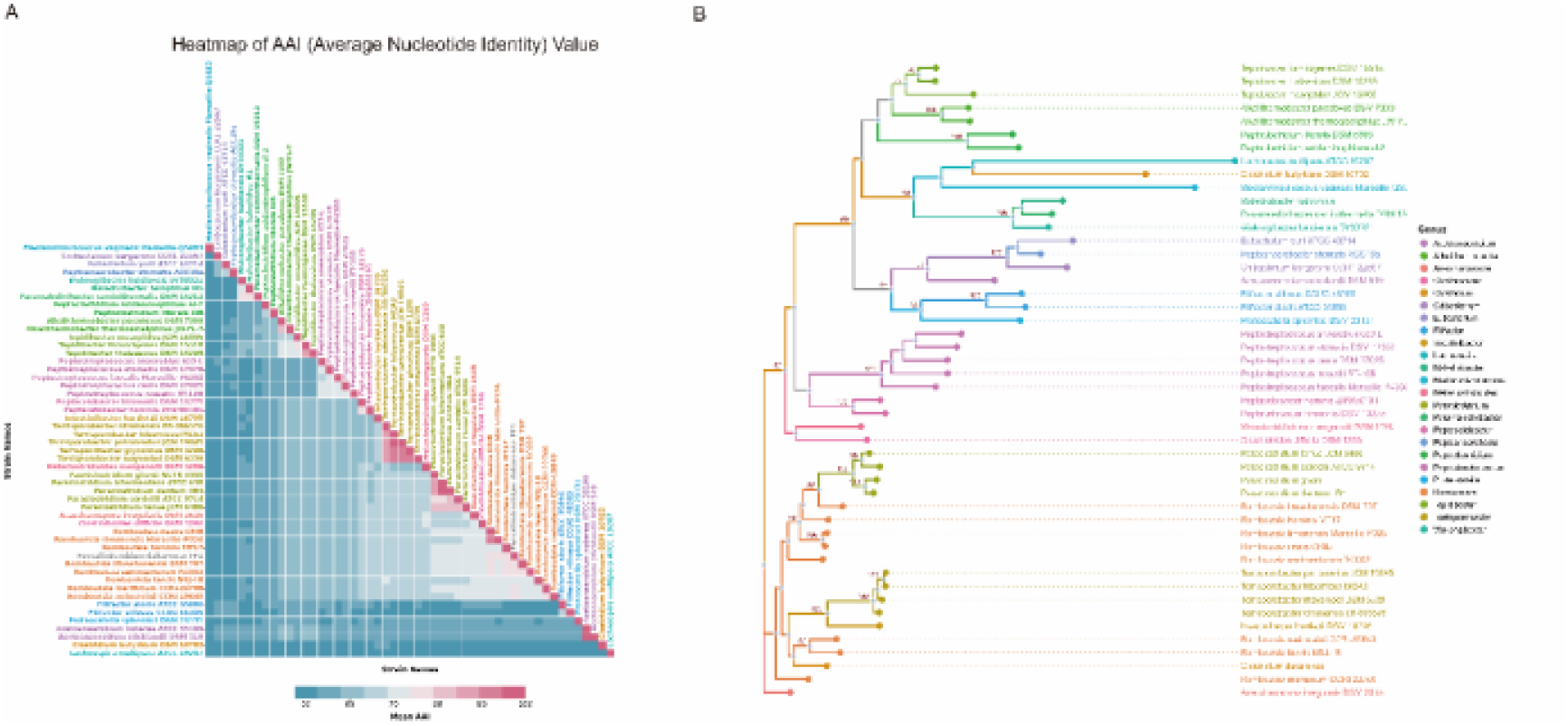
Amino acid identity analysis and phylogenetic tree based on core spore coat proteins

The phylogenetic conclusions based on spore coat proteins in this study align closely with those from AAI analysis and previous spore coat proteins and exosporium proteins phylogenetic studies, forming a multi-dimensional validation system that strengthens the reliability of taxonomic inferences. On the one hand, intra-genus AAI values are significantly higher than inter-genus values—for instance, the intra-genus AAI within *Terrisporobacter* is 90.68, which is higher than the low AAI values observed between distantly related genera (e.g., 79.17 between *Alkalithermobacter* and other genera, 58.96 in other cross-genus combinations). This AAI pattern corresponds to the topological differentiation of the spore coat protein tree, confirming that both molecular data types reflect consistent evolutionary relationships. On the other hand, spore coat proteins act as dual markers that simultaneously reflect evolutionary phylogenetic relationships (via sequence/clustering) and associations with spore formation phenotypes (via structural localization/function)—a unique advantage that enhances their taxonomic utility. Collectively, these multidimensional molecular indicators confirm that spore coat proteins are reliable markers for defining the taxonomic boundaries of the Peptostreptococcaceae family. They further provide systematic evidence for the precise revision of its taxonomic framework, including specific revisions supported by previous results: the merger of *Eubacterium* and *Paeniclostridium*, the establishment of a new genus for *Alkalithermobacter*, and adjustments to the taxonomic status of *Romboutsia* species (10). This further refines the classification and evolutionary research system of the family.

To further enhance the robustness of phylogenetic inferences, a maximum likelihood (ML) phylogenetic tree was constructed based on three conserved spore coat proteins (SpoIVA, CotE, YabG), which serve as key markers selected for their high conservation and functional relevance in spore assembly across *Peptostreptococcaceae*. These three core proteins were highlighted in the protein-protein interaction (PPI) network (Fig. 6) to visualize their functional connectivity, and this observation further supports their stability as taxonomic markers. Their consistent topological structures clearly represent the phylogenetic relationships among target groups (Figure 5B), providing direct evidence for the classification and delimitation of groups within the Peptostreptococcaceae family. In the trees, *C. difficile* forms a relatively independent monophyletic branch, with a significant genetic distance from adjacent groups (e.g., the genus *Peptoclostridium*), and it does not cluster with species from other genera—this topological feature emphasizes the phylogenetic uniqueness of *Clostridioides* as an independent taxonomic unit, strongly supporting the conclusion that it merits independent classification due to its molecular differentiation. Furthermore, within the genus *Romboutsia*, a clear phylogenetic differentiation pattern is observed—consistent with the earlier finding that *Romboutsia* is non-monophyletic: species such as *R. weinstockii* and *R. sedimentorum* cluster tightly, forming the genus’s core group, while *R. ilealis* and *R. timonensis* each form separate independent branches with considerable genetic distance from the core group. Notably, these two species do not form stable clusters with other closely related genera in the Peptostreptococcaceae family (e.g., *Paraclostridium*), supporting their classification as groups with uncertain taxonomic positions within the family and providing a topological basis for future genus-level revisions.

**Figure 6:**
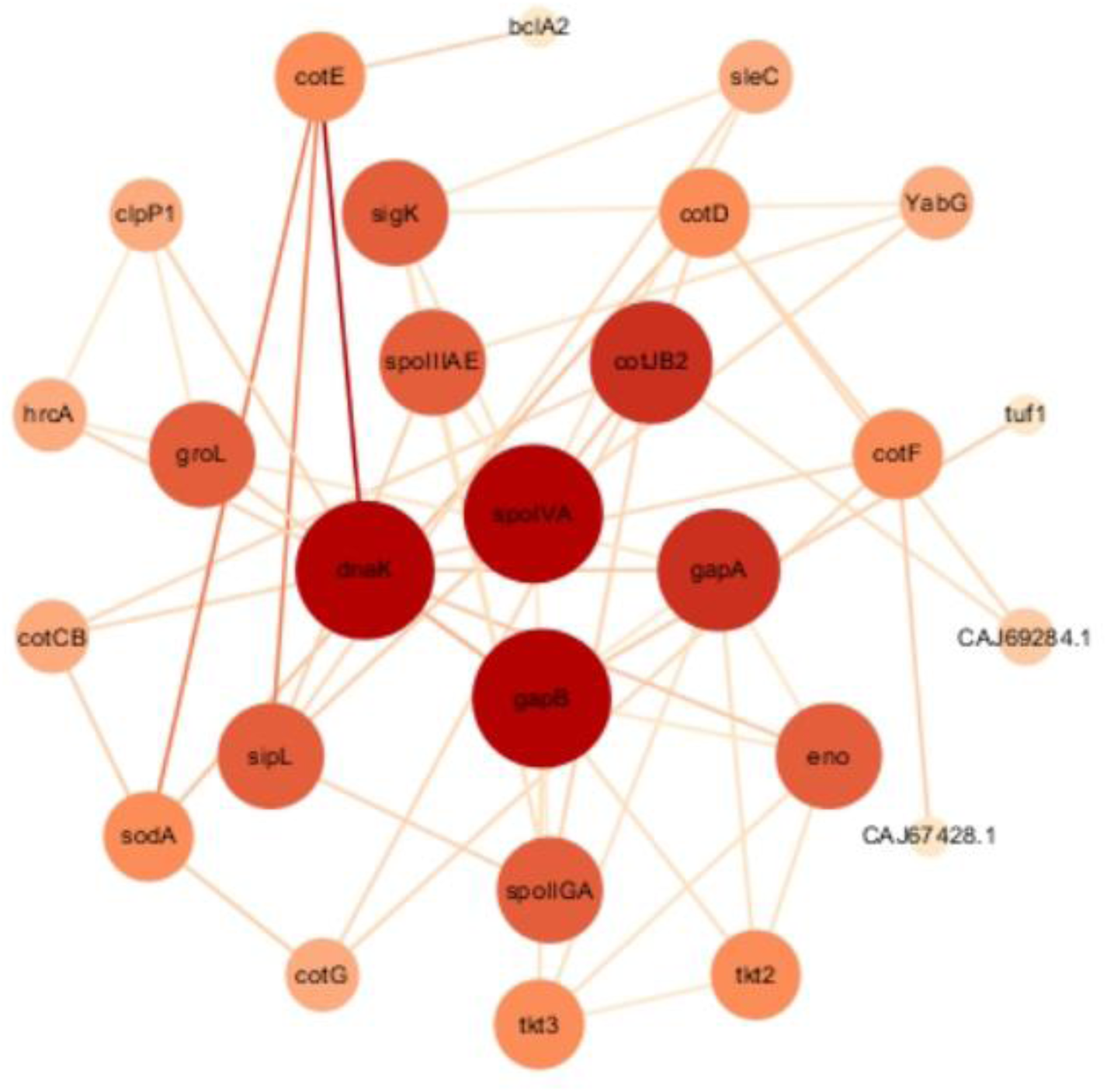
Spore coat/exosporium genes: Co-expression PPI network

Two key taxonomic revisions are supported by the ML tree results: first, species of the genus *Paraclostridium* (e.g., *P. ghonii*, *P. sordellii*) form closely clustered, high-support branches in the trees. Notably, *Eubacterium dentum* clusters with this genus’s core group, with Bootstrap support values ranging from 92% to 100%—this support is well above the 70% high-confidence threshold in phylogenetic analysis, directly validating the hypothesis of reclassifying *Eubacterium dentum* as *Paraclostridium dentum* and providing reliable phylogenetic evidence for this adjustment. Second, *Alkalithermobacter thermocaliphilus* and *A. paradoxus* form relatively independent monophyletic branches, clustering tightly within their respective branches and exhibiting significant genetic distances from other major clades in the Peptostreptococcaceae family (e.g., *Clostridioides* and *Romboutsia*). No cross-clade clustering is observed, supporting the conclusion that they should be classified into a new genus, *Alkalithermobacterium*, and providing crucial topological evidence for establishing this new genus.

### Analysis of Spore Core Proteins and Regulatory Mechanisms: Uncovering the Functional Basis of Conservation

This gene co-expression PPI network consists of 29 nodes and 98 edges (average degree 6.76). Circular nodes represent spore coat and exosporium genes. Arranged concentrically by connectivity degree, nodes have size and color positively correlating with degree (larger/darker = higher degree); genes with degree > average are defined as core targets. Node color intensity further reflects the co-expression Pearson correlation coefficient (PCC; darker = more significant), while edge thickness corresponds to the STRING combined score (0–1; thicker = more reliable interactions).

To explore interaction patterns among the identified spore coat proteins and exosporium proteins, we conducted protein-protein interaction (PPI) analysis using data from the STRING database. This analysis revealed 58 high-confidence interactions (edges) involving 27 genes (nodes), with an average connectivity degree of 3.87. The STRING combined scores for these interactions ranged from 0.700 to 0.999, and 65.5% of these interactions had a score of ≥0.8, indicating high reliability. Among the genes included in these high-confidence interactions, several core nodes exhibited notable connectivity, including the three conserved spore coat proteins (SpoIVA, CotE, YabG) highlighted in our phylogenetic analysis. Specifically, *spoIVA* (CD26290) had a connectivity degree of 7, interacting with *YabG* (CD35690), *sipL* (CD35670), and *sleC* (CD05510) (interaction scores: 0.728–0.912); *gapA* (CD31740) had a degree of 6; *eno* (CD31700) and *cotE* (CD14330) had degrees of 5 and 4, respectively; and both *clpP1* (CD33050) and *YabG* had a degree of 3. Notably, direct high-confidence interactions were observed between YabG and SpoIVA (score: 0.801) as well as between Eno and GapA (score: 0.987), with the former linking two of the three core spore coat proteins. Collectively, these interactions, particularly those centered on SpoIVA, CotE, and YabG, are likely pivotal for spore coat assembly as they integrate key structural and regulatory components of the assembly machinery.

To further define the functional relevance of these interactions, we classified genes with a connectivity degree exceeding the average (3.87) as core interaction targets, identifying 17 such genes (e.g., *dnaK*, *spoIVA*, *gapA*, *eno*, *cotE*). Importantly, the three core spore coat proteins (SpoIVA, CotE, YabG) are among these high-connectivity targets and exhibit the highest conservation within *Peptostreptococcaceae*, consistent with the sequence conservation patterns observed in Fig. 4A. While other core interaction targets display varying levels of conservation across the family, the exceptional conservation of SpoIVA, CotE, and YabG underscores their functional centrality in the spore coat assembly mechanism. This conserved interaction module, which is anchored by the three core proteins, further validates their suitability as robust taxonomic markers for resolving closely related species within *Peptostreptococcaceae*.

## Discussion

Conventional bacterial classification methods rely heavily on phenotypic characteristics, including morphological traits (e.g., cell size, shape, flagellation) (56), cultural properties (e.g., nutritional requirements, optimal growth temperatures), and biochemical test results. While intuitive, this approach is constrained by subjective interpretation and inherent technical limitations that compromise taxonomic accuracy (57). Modern classification has shifted toward genotypic data from DNA sequence analyses, yet 16S rRNA gene sequencing, still the most widely used single-gene marker, remains limited by gene recombination and mutation that obscure true evolutionary relationships and low sequence divergence among closely related taxa that restricts resolution and yields ambiguous assignments (58). To address these longstanding limitations, this study adopted a polyphasic taxonomic framework integrating the strengths of genome-scale precision and functional marker reliability. Specifically, we synthesized whole-genome single-copy orthologous protein sequence data, supplemented with average amino acid identity (AAI) calculations, and further validated these findings through cross-verification using spore proteins as taxonomic markers. This integrated strategy directly overcomes the shortcomings of traditional phenotypic approaches and single-locus genotypic methods, with whole-genome single-copy orthologous protein analysis as the core foundational component.

Phylogenetic analysis of whole-genome single-copy orthologous proteins has become a cornerstone of modern microbial taxonomy, with enhanced resolution for bacterial classification (59). These proteins are uniquely suited for phylogenetic inference due to their functional constraints, as they rarely undergo recombination, accumulate mutations at stable, clock-like rates, and retain sufficient divergence to distinguish closely related groups. Consistent with these inherent advantages, our whole-genome protein phylogeny (Fig. 2) clearly resolved core clustering patterns of key genera within Peptostreptococcaceae without conflicting signals. For instance, it confirmed the tight phylogenetic affinity between *Paraclostridium* and *Metaclostridioides*, the distinct evolutionary position of *Clostridioides*, and the putative non-monophyletic status of *Romboutsia*. These results align with recent taxonomic revisions based on comparative genomics (10), further validating the reliability of our foundational framework.

To further strengthen this framework, we integrated AAI calculations, a robust metric for quantifying genomic similarity that independently validates taxonomic boundaries (60). AAI is widely adopted for genus-level classification, with a threshold of ≥65–70% typically defining members of the same genus (61). Our AAI data strongly corroborated phylogenetic insights from the single-copy protein tree, as the mean AAI of 58.96 between *Alkalithermobacter* and *Peptacetobacter* fell well below this threshold, reinforcing their distinct generic status. Additionally, genetic distance data from the whole-genome protein tree showed the distance between the *Clostridioides difficile* DSM 1296 clade and the *Metaclostridioides* clade is approximately 0.9, far exceeding the typical intra-genus divergence threshold (≤0.05 for most bacterial genera). This convergence of phylogenetic topology, genetic distance, and AAI evidence, supported by extensive prior literature, underscores the rigor and robustness of our whole-genome-based foundational framework.

Building on this reliable whole-genome framework, we further integrated conserved spore coat/exosporium protein sequences into the polyphasic taxonomic strategy. Notably, our findings regarding *Terrisporobacter* are highly consistent with prior studies indicating this genus belongs to the Peptostreptococcaceae sensu stricto monophyletic clade (10). This consistency confirms *Terrisporobacter*’s core taxonomic position within this family and reinforces the reliability of our integrated framework.

Employing spore coat proteins as specialized molecular markers yielded critical taxonomic insights complementing genome-scale analyses. For instance, it accurately assigned *C. difficile* to the genus *Clostridioides* and phylogenetically distinguished the non-pathogenic *Metaclostridioides mangenotii* from this taxon. The molecular divergence observed in spore coat proteins provides direct evidence for *Clostridioides*’ taxonomic distinctiveness from its close relatives, supporting its status as a phylogenetically robust independent unit. Intergeneric genetic differences revealed by AAI and whole-genome analyses further corroborate clear genetic boundaries within Peptostreptococcaceae, aligning with inferences from spore protein data. Together, these lines of evidence, including spore coat protein divergence and congruent genome-scale genetic signals, validate the phylogenetic independence of *Clostridioides*, consistent with prior observations that *C. difficile* harbors spore-associated sequence specificity driven by unique genetic events.

The conserved nature of spore coat proteins, a key factor in their taxonomic utility, stems from functional constraints imposed by their role in spore assembly and survival. Specifically, spore coat assembly relies on specific protein-protein interactions (PPIs), so a single protein cannot evolve independently without disrupting its obligate partners, thereby reducing mutation likelihood. This constraint contrasts sharply with *C. difficile* toxins such as TcdA and TcdB, which evolve in response to environmental factors like host microenvironments and exhibit high variability via recombination (62), highlighting the superiority of spore coat proteins as stable taxonomic markers. Furthermore, the spore coat’s function as a barrier against enzymatic degradation and antibiotics (63) is thought to be linked to its assembly, which may be controlled by a protein homeostasis network previously shown to maintain protein integrity in related systems.

This conserved assembly process is regulated by a protein homeostasis network centered on the *dnaK-groL* chaperone system and the *clpP1* protease. For instance, *clpP1* interacts with both *dnaK* and *groL*, with interaction scores ranging from 0.775 to 0.875. This regulatory network ensures proper folding and quality control of spore coat proteins, as supported by previous studies (64–66). Phylogenetic analyses based on spore coat proteins, together with 16S rRNA sequencing, AAI calculations, and ML/NJ tree constructions, established consistent taxonomic signals validating the phylogenetic utility of spore coat proteins. Meanwhile, high-confidence PPI patterns, conserved core targets, and the underlying regulatory network provided a mechanistic basis for their sequence conservation, a key prerequisite for reliable taxonomic markers. Together, these complementary lines of evidence, namely systematic phylogenetic congruence and robust functional and network conservation, strongly support establishing spore coat proteins as a novel molecular marker for Peptostreptococcaceae taxonomy.

Notably, taxonomic inferences from spore protein-based phylogenies are highly congruent with those derived from the whole-genome protein phylogeny (Fig. 2), with no conflicting signals in the core clustering patterns of key genera. This cross-validation between two distinct marker systems, spore-specific proteins and whole-genome proteins, directly corroborates the reliability of spore protein-based classification inferences. Furthermore, the spore protein-based NJ tree exhibited robust bootstrap support (≥70%) for core branches, and genetic distance data from whole-genome analyses further reinforced the tree’s ability to resolve closely related genera. Collectively, the high congruence between spore protein and whole-genome protein phylogenies, together with corroborative genetic distance evidence from Fig. 2, unequivocally demonstrates the feasibility of spore-associated proteins as credible taxonomic markers for Peptostreptococcaceae and related spore-forming bacteria.

Our study also directly addresses unresolved questions in Peptostreptococcaceae taxonomy and spore biology. Specifically, our comparative analysis identified 11 highly conserved proteins across 51 bacterial species, an area hindered by unknown proteins and unclear layer composition (67). This set may include uncharacterized morphogenetic proteins and provides valuable targets for future spore assembly studies. To enhance genus-and species-level classification, we propose exploring additional core conserved proteins such as single-copy housekeeping proteins with stable evolutionary rates. This expansion will address limitations of single markers such as 16S rRNA genes and support efforts to revise *Romboutsia*’s generic boundaries and verify *Terrisporobacter*’s taxonomic status.

In summary, our polyphasic taxonomic framework, anchored in the rigor of whole-genome single-copy orthologous protein phylogeny and AAI calculations and augmented by the novel application of spore coat proteins, overcomes limitations of traditional nucleic acid methods by leveraging a three-dimensional framework encompassing structural conservatism, functional relevance, and ecological adaptability, with each dimension supported by our integrated data. Practically, this approach has resolved genus-level controversies surrounding *C. difficile* and confirmed *M. mangenotii*’s independent evolutionary status, while paving new avenues for taxonomic studies of spore-forming genera such as *Bacillus* and *Clostridioides* and refining microbial classification systems within the phylum Firmicutes. Furthermore, our results are consistent with prior conclusions from concatenated core protein trees, validating that functionally conserved spore coat proteins ensure phylogenetic stability and reinforce taxonomic placements within Peptostreptococcaceae.

